# VenomKB v2.0: A knowledge repository for computational toxinology

**DOI:** 10.1101/295204

**Authors:** Joseph D. Romano, Victor Nwankwo, Nicholas P. Tatonetti

**Affiliations:** Department of Biomedical Informatics, Columbia University, New York, NY, 10032.; Department of Systems Biology, Columbia University, New York, NY, 10032.; Institute for Genomic Medicine, Columbia University, New York, NY, 10032.; Department of Medicine, Rush University Medical Center, Chicago, IL, 60612.; Department of Medicine, Columbia University, New York, NY, 10032.; Data Science Institute, Columbia University, New York, NY, 10032.

## Abstract

**Motivation:** Venom peptides comprise one of the richest sources of bioactive compounds available for drug discovery. However, venom data and knowledge are fragmentary and poorly structured, and fail to capitalize on the important characteristics of venoms that make them so interesting to the biomedical community.

**Results:** We present VenomKB v2.0, a new open-access resource for knowledge representation and retrieval of venom bioactivities, sequences, structures, and classifications. VenomKB provides a complete infrastructure for computational toxinology, with a focus on drug discovery and effects that venoms have on the human body. VenomKB is accompanied by a suite of tools for programmatic access, and, in this article, we highlight scenarios demonstrating its usefulness and novel contributions to toxinology, pharmacology, and informatics.

**Availability:** VenomKB can be accessed online at http://venomkb.org/, and the code can be found at https://github.com/tatonetti-lab/venomkb/. All code and data are available under open-source and open-access licenses.

## 1 Introduction

Animal venoms are complex mixtures of proteins, carbohydrates, and other compounds that are used for both defense and predation in millions of species distributed widely across the tree of life [Calvete et al., 2009, Kaas and Craik, 2015]. Due to their immense combinatorial diversity and strong, targeted effects in biological systems, venom proteins are of considerable interest to the drug discovery and biotechnology communities [Harvey, 2014]. In the United States, approximately 20 venom-derived drugs have been approved by the Food and Drug Administration to-date, with many more currently undergoing clinical trials [Lewis and Garcia, 2003, Mafong and Henry, 2008, Sanford, 2013].

Toxinology—the study of venoms and other naturally-occurring toxins—is a highly active scientific discipline. Toxinologists characterize the individual proteins found in venoms using a combination of transcriptomics, proteomics, and traditional biological assays [Calvete et al., 2009, Gon¸calves-Machado et al., 2016]. Further, venom components are associated with clinically important diseases and conditions (in a previous study, we found almost 40,000 unique mentions of venoms treating a disease in MEDLINE alone [Romano and Tatonetti, 2015]). Despite these areas of rapid scientific productivity, there is an unmet need for a resource that aggregates and structures these relatively disparate types of information and knowledge [Jensen et al., 2014, Paul et al., 2010, Wishart et al., 2017].

Here, we present VenomKB v2.0—a new resource for aggregating and representing venom knowledge including molecular characteristics, biodiversity data, manually- and automatically-identified literature data, and a standardized ontological representation for these different data types. VenomKB v2.0 is a complete rewrite of a previous toxinology resource aimed specifically at literature data [Romano and Tatonetti, 2015], the contents of which are included in v2.0 in a more controlled and robust format. VenomKB is built with a modern and intuitive interface along with a REST API to make all data elements programmatically available. This knowledge base is the most complete public resource for computational toxinology research to-date, and it stands to become a major resource for toxinologists, informaticians, molecular biologists, and educators interested in venoms and/or their components.

## 2 Results

VenomKB can be accessed online at http://venomkb.org/. The original version of the knowledge base is still available for use, and can be accessed via a link on the home page of the URL above.

### 2.1 Size and structure of VenomKB

VenomKB currently catalogues 6,236 venom proteins from 632 venomous species of animals. VenomKB also contains five genomes from venomous animals, which—at the time of writing— is the entirety of publicly available venomous animal genomes known to the authors. The major data types in the knowledge base are summarized in **Table 1**. **Figure 2** shows counts of the various data types contained in VenomKB.

**Table 1:** VenomKB size and data types

| Data type | Number of Records | VenomKB ID Prefix |
| --- | --- | --- |
| Proteins | 6,236 | P |
| Species | 632 | S |
| Genomes | 5 | G |
| Disease/condition annotations | 1,065 | E |
| Literature predications | 14,710 | – |
| Gene Ontology annotations | 18,677 | – |

**Figure 1:**
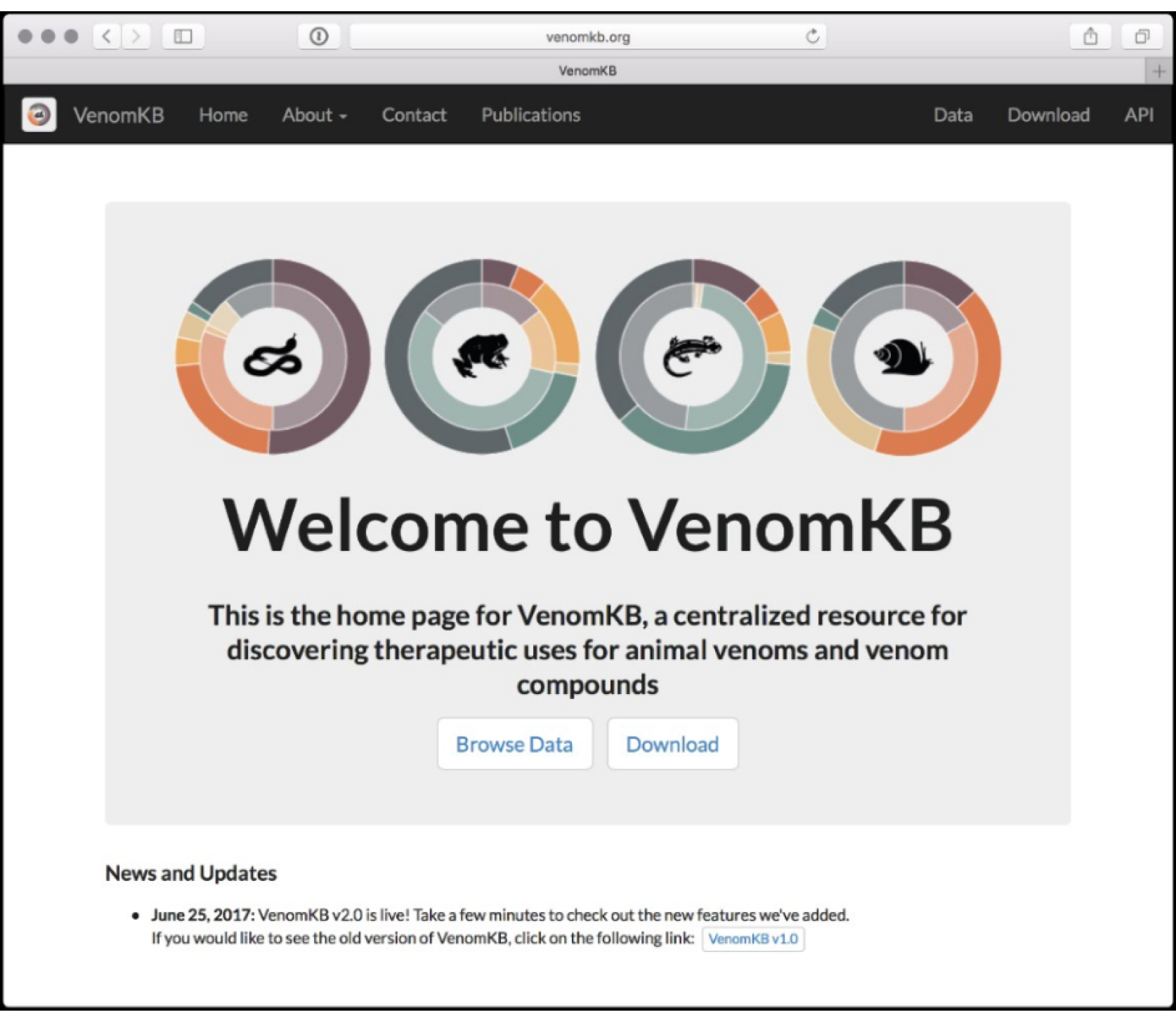
Image of the home page for VenomKB. Users can access data and informational pages via the navigation bar or in the main body of the website. A “News and Updates” section provides useful information and changes made to the website.

**Figure 2:**
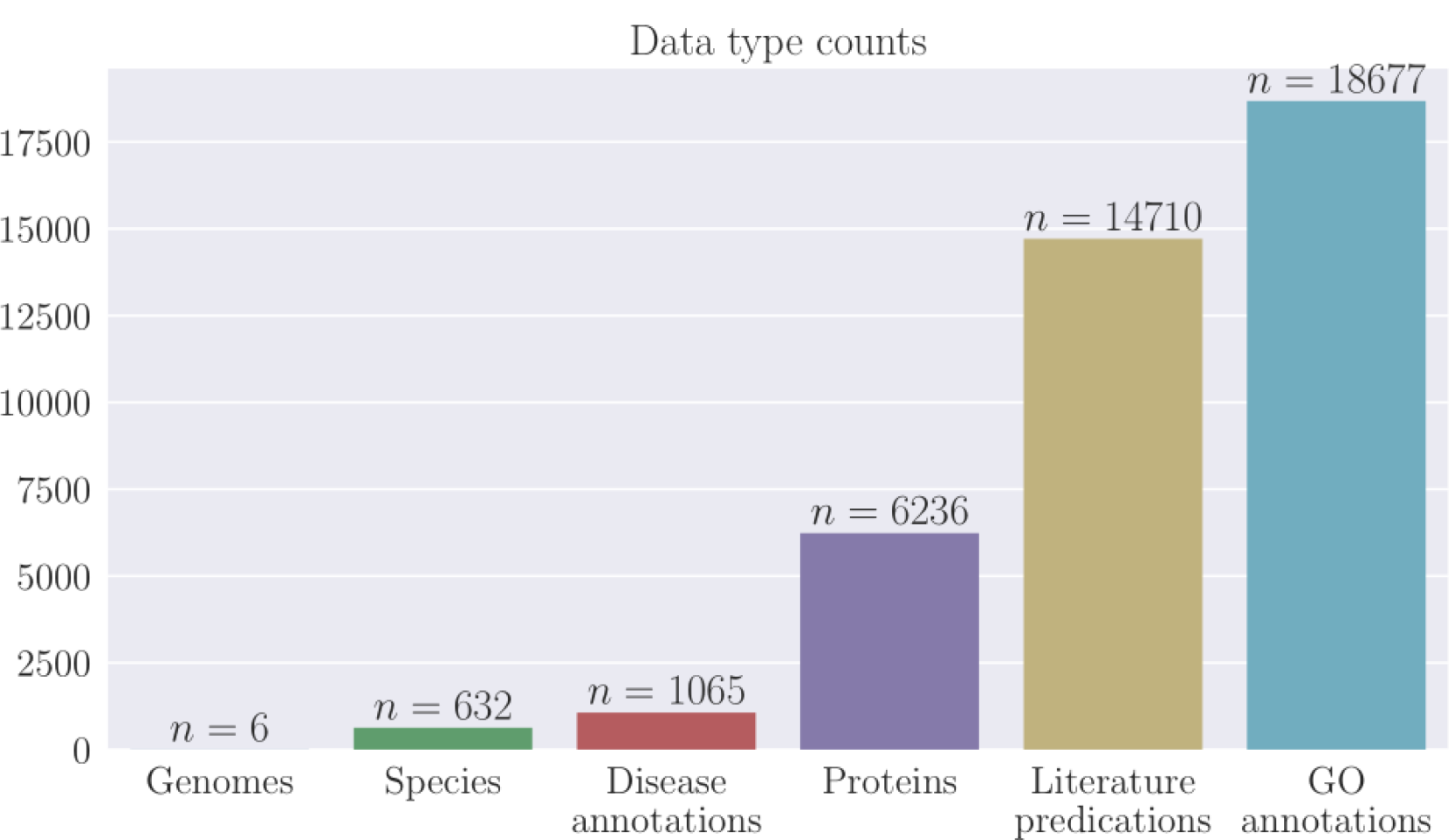
Barplot of counts of data types in VenomKB v2.0. Genomes, Species, and Proteins are ‘primary’ data types represented as instances of Venom Ontology classes; Disease annotations, Literature predications, and GO annotations are ‘secondary’ data types that are represented as properties of primary data types.

Each of the previously described data types is structured according to the Venom Ontology [Romano and Tatonetti, 2016], which provides a formal description of the different types of data related to venoms, along with the types of relationships that exist between them. Every data record in VenomKB is assigned a unique, permanent identifier that consists of one alphabetical character followed by seven digits. The first character indicates the data type (see **Table 1**), and the seven digits are randomly assigned.

We sourced all non-inferred data in VenomKB from other publicly available resources. A large number of the molecular/protein data were adapted from UniProtKB/Swiss-Prot’s Tox-Prot annotation system [Jungo et al., 2012], which is a major effort to identify and manually curate animal toxin peptides (including venom components) in UniProtKB. The number of proteins (6,236) currently in VenomKB is equal to the number of venom components in Tox-Prot at the time of constructing the database.

VenomKB also contains 39,179 literature annotations that describe a venom or a venom component treating a disease or health condition, which we transferred from VenomKB v1.0. Of these, 275 were manually curated, and 33,284 are normalized semantic predications extracted from the Semantic MEDLINE database using a knowledge discovery approach, which is described in a previous study [Romano and Tatonetti, 2015]. We automatically mapped 14,710 of these predications to both species and individual proteins using ontological inference; these predications are shown in both the Species and Protein data pages, as well as the raw JSON representations of these data types.

### 2.2 Web application description

The home page for VenomKB is shown in **Figure 1**. From the home page, users can access most components of the web application, as well as a link to the VenomKB v1.0 application, for backwards compatibility. The main interface for exploring data is located at http://venomkb.org/data, or from links on the home page. The interface is shown in **Figure 4**, for reference. Users can filter data records in several ways, including by name, data type (e.g., proteins, species, or genomes), annotation score (1 to 5 stars, explained below), or by disease/condition annotations. Data types not included in this interface (such as literature predications and other annotations) are embedded within the structure of their corresponding documents. The search interface allows sorting by column.

When users find a data record of interest, they can view it by clicking on its corresponding VenomKB ID (VKBID), or by navigating to ‘http://venomkb.org/{VKBID}’. An image of a protein detail page is shown in **Figure 5**. The detail page for individual data records presents information that is not available in the data search interface (e.g., for proteins, this includes amino acid sequence information, Gene Ontology annotations, literature predications, related articles from PubMed, a link to the species the venom is from, and others). Furthermore, tabs at the top of the data detail page allow the user to view the record in JSON (JavaScript Object Notation) format or download the record as a JSON text file. Users can run BLAST on the amino acid sequences for protein data records, and we plan to add other external analysis tools in the near future. Whenever possible, species pages provide a complete taxonomic lineage for the venomous species being described (the major exception to this is for some species of scorpion, which are interestingly underrepresented in ITIS—the public database we used to source taxonomies). Where appropriate, an image is displayed showing the current data element. At the bottom of each page is a list of external identifiers corresponding to the element currently being viewed. If users find an error in any given data element, a button allows them to report the issue to the website’s administrator.

**Figure 3:**
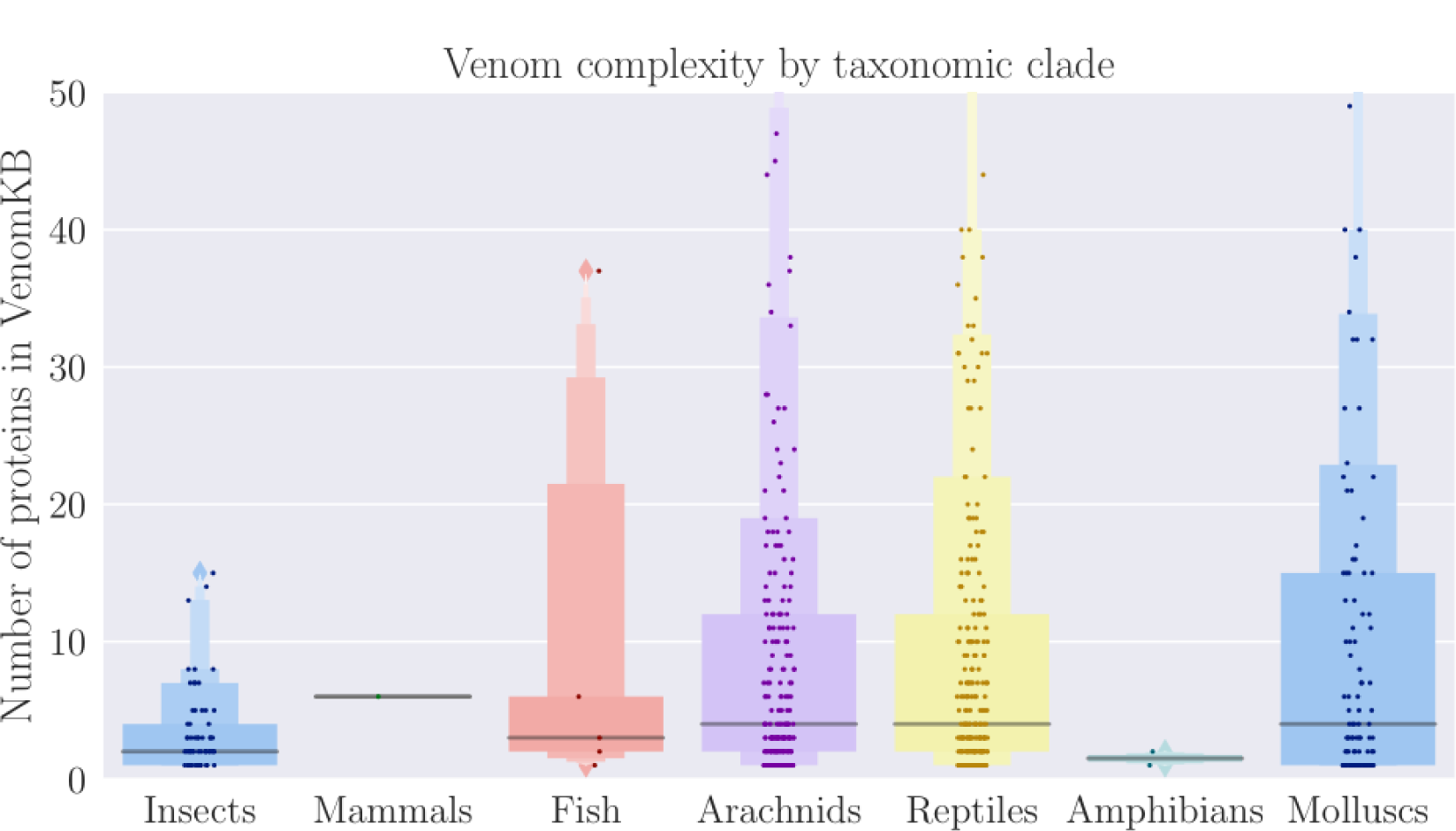
Venom complexity by major taxonomic groups. ‘Complexity’ is defined as the number of proteins present in VenomKB for a specific venomous species. The relatively low complexity of insect venoms compared to arachnids, reptiles, and molluscs could be informative for the purposes of drug discovery.

**Figure 4:**
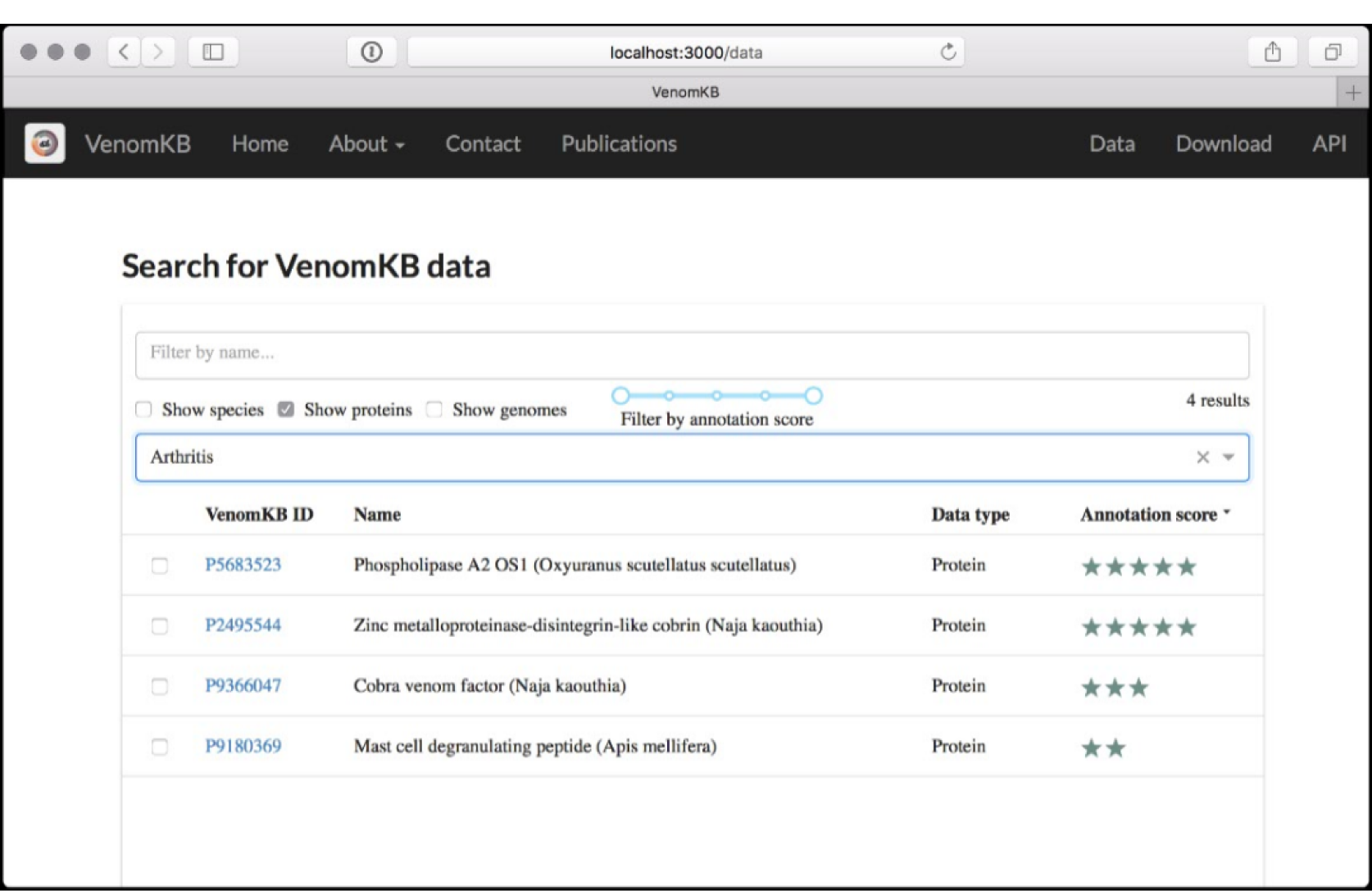
Interface for graphically searching and browsing data in VenomKB. Users can search by string, data type, data quality score, and by disease/condition annotation. The query results page allows sorting by various fields. To access a particular data record, click on the VenomKB ID corresponding to the entry of interest.

**Figure 5:**
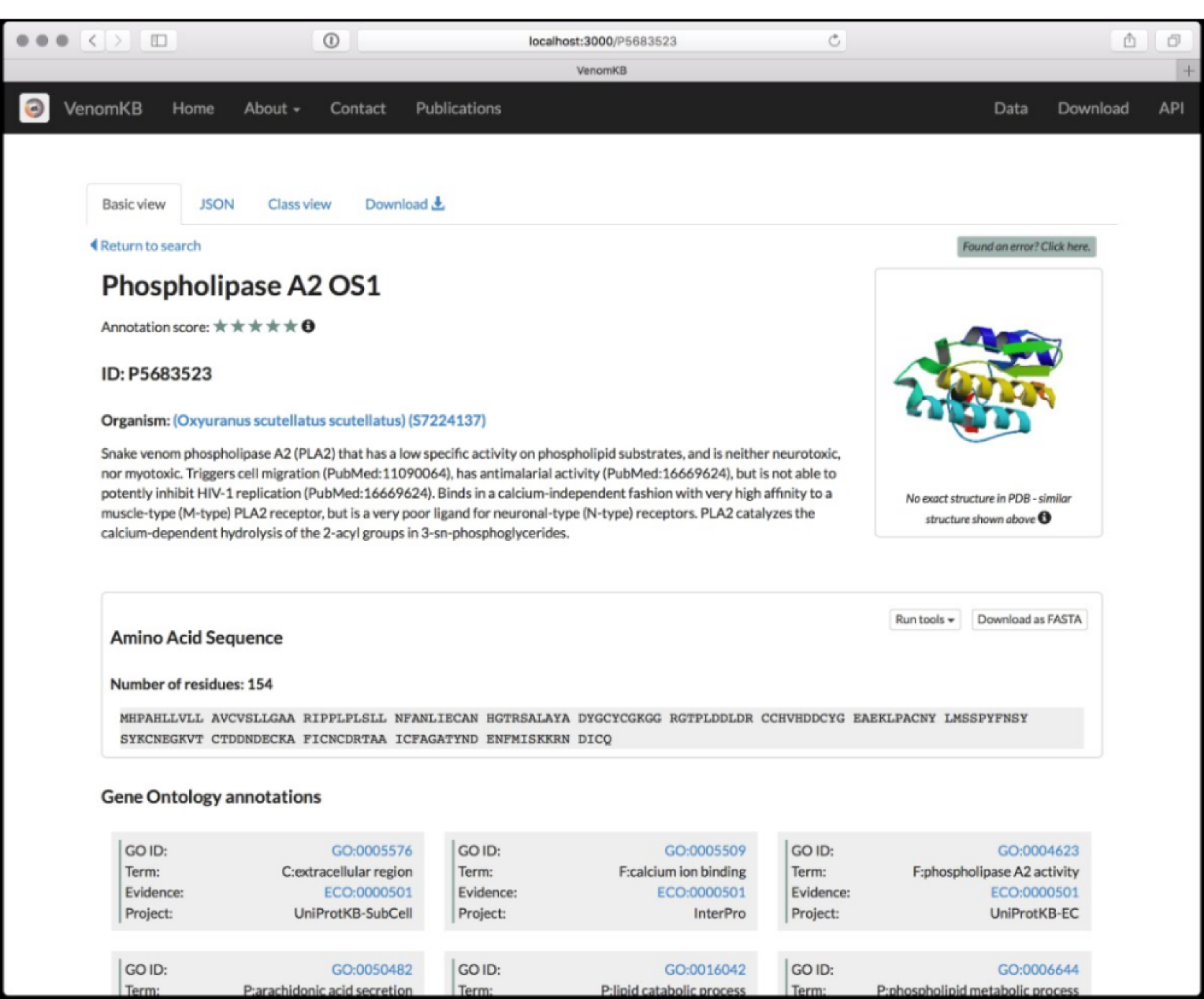
Example of a page containing a single venom protein. Userschoose the way that they view data using the tabs at the top of the interface. The user is presented with an image of the protein, descriptive information, a link to the species from which the protein was discovered, amino acid data (with links to external tools such as BLAST), and gene ontology annotations. Other fields are out of view, including literature predications, links to external databases, and related publications from MEDLINE.

### 2.3 VenomKB augments existing knowledge using ontological inference

There are generally two types of ontological inference in VenomKB, both of which are dependent on the structure of the Venom Ontology: 1.) Inferred data types and 2.) inferred data associations. Currently, the only inferred data type in VenomKB is “Systemic Effects”, which are diseases and conditions that are either associated with or resulting from the administration of a venom or venom component to the human body. Another inferred data type that we plan to add in the future is “Molecular Effects”, which are the specific effects that venoms and their components have on biomolecular structures at the cellular or sub-cellular level in the human body. By using the structure of the Venom Ontology, we can use class assertions to infer and validate the molecular effects associated with diseases and conditions, and potentially discover new disease/condition associations for venoms and venom components.

In Figure 3 we illustrate a specific example of the type of observation that can be made using a combination of VenomKB’s data and ontological inference. Here, we define venom complexity as the number of unique protein components in a species’ venom. By grouping species in VenomKB using the available taxonomic hierarchy and then counting the number of linked protein records for those species, we can plot distributions of venom complexity for major taxonomic clades, such as reptiles, insects, molluscs, and others. In addition to highlighting the relative lack of mammals, fish, and amphibians in VenomKB (and, by extension, other databases containing venom data), these distributions highlight that insect venoms seem to be of lower complexity than arachnid, reptile, and mollusc venoms. This observation may be useful for the purposes of drug discovery—for example, it could suggest that components of insect venom tend to be less specific in their molecular targets, perhaps so they have activity in a wider range of species (which is well-supported in the literature of evolutionary toxinology) [Schmidt, 1990, Kordiš and Gubenšek, 2000].

### 2.4 Heuristic annotation scores provide a relative measure of data quality

A major aspect of creating publicly available databases for science is to provide methods for assessing the quality of the data. Data quality can be assessed using two general approaches: task-based assessment, and by performing intrinsic tests on the data records. Intrinsic assessments of data quality are challenging, especially when designing inferred data types that lack a baseline reference. One noteworthy example of addressing this issue is in the UniProt database, where data elements are given scores that indicate completeness and confidence in the assertions made by that element. However, few structured databases outline an objective approach to assigning quality scores.

We defined heuristic annotation scores for each data record in VenomKB, which are designed to provide a means for comparing the quality of VenomKB data entities relative to all other entities of the same data type. These scores are represented as integers in the range [1 ‥ 5], inclusive, and are displayed as ‘star’ icons on the data browse and data detail pages in the web application. To ensure that these quality measures are well-distributed within each data type, we balanced the number of elements attaining each of the five possible scores. The procedure we used to create these annotation scores is described in Experimental Procedures.

### 2.5 Data availability and programmatic access to VenomKB

All data and code related to VenomKB are freely and publicly available online. A version-controlled Git repository for a.) generating the database back-end and b.) the VenomKB web application itself can be accessed at http://github.com/jdromano2/venomkb. The code used to generate the database is written in the Python programming language, and it uses the PyMongo library to populate a MongoDB database instance with the generated data. The web application is written in JavaScript (using the React library to design the user interface and Redux to represent the internal state of the data model), and communicates with the Mon-goDB back-end via a REST API (Application Programming Interface) that is also accessible for programmatic access by end-users. The API functionality is documented on VenomKB’s website at http://venomkb.org/about/api.

## 3 Discussion

### 3.2 Advantages of VenomKB over existing venom databases

To our knowledge, VenomKB is one of five public databases focused on venoms and their components. In designing VenomKB, we aimed to improve on a number of characteristics that make these databases unsuitable for many tasks. The other four databases are UniProtKB/Swiss-Prot’s Tox-Prot dataset [Jungo et al., 2012], the University of Queensland’s ConoServer [Kaas et al., 2010, Kaas et al., 2011] and ArachnoServer [Pineda et al., 2017] databases, and the Animal Toxin DataBase (ATDB) [He et al., 2010]. ConoServer and ArachnoServer are each focused on specific clades of venomous animals (cone snails and arachnids, respectively). Tox-Prot is a relatively small component of the much larger UniProtKB, and therefore does not have the ability to support many of the characteristics unique to venoms. ATDB seems to no longer be available for public use, at the time of writing.

VenomKB seeks to address each of these shortcomings. Of particularly critical importance is VenomKB’s inclusion of several datatypes that are present in none of the alternative venom databases. This includes inferred disease/condition associations, explicit representations of the animal species from which the proteins are derived, publicly available genome data, and the semantic predications extracted from previous scientific publications. As described by [Frijters et al., 2010], types of data like these are critical to the drug discovery process. For example, if a protein has a known therapeutic effect but is too toxic to administer to humans, similar species may synthesize less toxic alternatives.

**Table 2:** Version differences; VenomKB v1.0 vs. v2.0

|  | **VenomKB v1.0** | **VenomKB v2.0** |
| --- | --- | --- |
| Web framework | Ruby on Rails | Node.js + React + Express |
| Database back-end | PostgreSQL | MongoDB |
| Database structure | 3 unstructured SQL tables | Structured documents mapped to Venom Ontology |
| API | None | REST API, implemented in Mongoose.js |
| Legacy support | n/a | VenomKB v1.0 rows mapped to v2.0 documents |

**Table 3:** Data sources

| Data source | Used for |
| --- | --- |
| ToxProt | Most molecular data in “Protein” records |
| NCBI Taxonomy | Species nomenclature data |
| ITIS | Species taxonomic hierarchies |
| Protein Databank (PDB) | Protein images |
| Wikimedia Commons | Species images |
| MEDLINE / SemMedDB | Structured semantic predications |
| VenomKB v1.0 | Raw semantic predications |
| Gene Ontology | GO protein annotations |

VenomKB is not limited to certain clades of venomous species. In addition to improving the coverage of the data, this also allows users to compare characteristics of venoms that have similar properties despite coming from unrelated species. However, it does limit its focus to venoms and concepts related to venoms, which allowed us to structure the knowledge base around the Venom Ontology and exploit the unique semantic features of venoms in a way to make inferences that would otherwise be challenging. We specifically host VenomKB on its own domain (venomkb.org) instead of on an institutional website: Since institutional websites and affiliations tend to change, having a dedicated domain name improves the site’s sustainability model.

### 3.2 Extending the VenomKB technique beyond venoms

Although VenomKB was designed specifically to manage venoms and venom component data, it is reasonable to assume that our techniques could be extended to other similar domains of interest. Plant metabolites, in particular, provide an interesting target, especially given that they already comprise a major source of approved therapeutics worldwide [Demain, 2002, Ngo et al., 2013]. The process of translating the structure of VenomKB to another domain would essentially involve three steps: (1) redefining the ontology on which the knowledge base is built (e.g., creating an appropriate plant metabolite ontology), (2) finding the appropriate data sources for populating the knowledge base, and (3) making inferences to define new data types where possible.

### 3.3 VenomKB as a model for open access of scientific data

As mentioned in the previous section, all code and data related to VenomKB are freely accessible to the public. These resources are maintained under the open-source GNU General Public License v3, which permits use, reuse, and modification under limited terms. A copy of this license is distributed as part of the source code repository.

### 3.4 Limitations

VenomKB is limited by a general lack of availability of venom data. Given that scientists believe there may be millions of venomous species on the planet [Smith and Wheeler, 2006], the 632 species represented in VenomKB comprise only a miniscule fraction of the total. This disparity is even more apparent when viewed from the perspective of whole-genome sequencing data: VenomKB only contains 5 species’ whole genomes (which, as stated before, is the entirety of publicly available genomes from venomous species, at the time of writing). This issue is exacerbated further by the fact that it is often challenging to tell whether a species is venomous or not—for example, it was only discovered in 2009 that the common octopus (*Octopus vulgaris*) is venomous, since the octopus is neither aggressive, nor is the venom appreciably toxic to humans [Ruder et al., 2013].

Although VenomKB contains novel data in the form of literature predications and automatically inferred disease/condition associations (as well as the ontological relationships between datatypes), much of the knowledge base is aggregated from previously compiled data sources, such as UniProtKB, NCBI, and others. However, in the near future, VenomKB will soon include novel experimental data in the form of human gene expression profiles that capture transcriptional responses to being exposed to specific animal venoms.

### 3.5 Future additions to VenomKB

VenomKB is—and likely will remain—a work in progress. Our goal is to provide a *comprehensive* knowledge resource for computational toxinology, but due to both the breadth of venom data types (experimental, clinical, molecular, etc.), and the rapid generation of new venom data, it is unlikely that any venom data resource will ever be truly comprehensive.

To address this challenge, a crucial aspect of VenomKB is a map of current and planned features that grows with and adapts to the evolving needs of the toxinology and drug discovery communities. This feature map is available to view at http://venomkb.org/about/features/. Aside from the novel gene expression profile data that was mentioned previously, important additions in the near future include the following:

- Important pharmacokinetic and biochemical measures (when known), such as IC_50_, *K_i_*, and molecular mass
- Additional gene-level data, including nucleotide sequences, protein isoforms, and gene families
- Annotations to clinical trials exploring particular venom compounds
- Metrics related to whole genomes, such as total size and sequencing methods used
- Species-level data related to natural uses of venoms, such as predation/defense, venom delivery, and target species

Furthermore, we strongly encourage input from researchers who could benefit from additional features. Contact methods for the authors are provided on the VenomKB website, at http://venomkb.org/contact.

## 4. Methods

The original version of VenomKB was written using the Ruby on Rails web framework for the Ruby programming language, but for v2.0 we rewrote the entire web application in JavaScript, using the React.js library to implement the interactive user-interface, and the Mongoose library to construct the data model for the REST API. The differences between v1.0 and v2.0 are summarized in **Table 2**. We maintain the database back-end for VenomKB on a MongoDB server that is separate from the web application for security and performance.

We constructed the database using an iterative approach, starting with data aggregated from existing databases and then transitioning to the addition of inferred and novel data types. To serve as a starting point for building the database, we treat the ToxProt venom protein annotation program as a gold-standard, being arguably the most complete existing venom database that is not constrained to a certain set of taxa. First, we retrieved all venom peptides in the ToxProt database and extracted core attributes relevant to VenomKB (such as amino acid sequences and cross-references to other databases). We then retrieved taxonomy data for all species with at least one peptide, and used both the NCBI Taxonomy database [Federhen, 2011] and the Integrated Taxonomic Information System (ITIS) to build taxonomic lineages and to retrieve other species-level data, such as common names, synonyms, and external identifiers.

To link literature annotations and predication data from VenomKB v1.0 to the new knowledge base, we used expert-identified literature references provided by the ToxProt program. For each PubMed identifier in ToxProt, we retrieved corresponding VenomKB v1.0 predications, and linked them to both their respective protein and species data records. Since many literature annotations are duplicated both within a single document and between multiple documents, we merged duplicate records.

In VenomKB, we represented data provenance using the PROV-DM data model standard [Missier et al., 2013]. Data provenance is a representation of the sources of each data type in VenomKB along with the methods employed to manipulate and restructure data. Beyond accountability and reproducibility, provenance allows for data quality assessment [Hartig and Zhao, 2009]—data aggregated, created, or validated by more rigorous methods generally are deemed to be of better quality than otherwise. The provenance model for VenomKB can be downloaded from the website, at http://venomkb.org/download.

### 4.1 Generating balanced heuristic annotation scores

In Results we explain the use of heuristic annotation scores to provide a method for comparing data quality and completeness relative to VenomKB’s other data elements of the same type. We accomplished this task by first assigning raw (unscaled) scores to each instance of each data type based on presence and absence of certain elements. For example, the raw score of a protein was increased by 0.05 for each literature predication, and decreased by 0.2 if it had no literature predications. A species’ raw score was increased by 3.2 if a complete taxonomic lineage was present, and decreased by 1.0 if no image of that species was available. The complete details for assigning raw scores is outlined in the VenomKB code repository. After computing raw scores, we then adjusted the scores for each data type to a discrete uniform distribution on the range [1..5] using the following transformation:

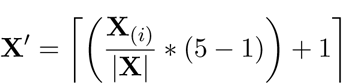

where *|***X***|* denotes the number of elements of data type **X**, **X**_(*i*)_ is the vector of order statistics for the raw scores of data type **X**, and **X^′^** is the vector of transformed scores. This procedure produces five evenly sized bins from 1 to 5 for each data type in VenomKB.

## Acknowledgements

We would like to thank Chunhua Weng and Adler Perotte for valuable comments and suggestions during the preparation of VenomKB.

## Funding

This work was supported by the National Institute for General Medical Sciences (grant R01 GM107145. PI: Tatonetti) and the National Center for Advancing Translational Sciences (grant OT3 TR002027. PIs: Tatonetti, Dumontier, and Weng).

